# Hypoxia-inducible factor 2α promotes protective Th2 cell responses during intestinal helminth infection

**DOI:** 10.1101/2025.01.09.631414

**Authors:** Jasmine C. Labuda, Tayla M. Olsen, Sheenam Verma, Samantha Kimmel, Thomas H. Edwards, Matthew J. Dufort, Oliver J. Harrison

## Abstract

Th2 cells must sense and adapt to the tissue milieu in order to provide protective host immunity and tissue repair. Here, we examined the mechanisms promoting Th2 cell differentiation and function within the small intestinal lamina propria. Single cell RNA-seq analyses of CD4^+^ T cells from the small intestinal lamina propria of helminth infected mice revealed high expression of the gene *Epas1*, encoding the transcription factor hypoxia-inducible factor 2a (HIF2α). *In vitro*, exposure to hypoxia or genetic HIF2α activation promoted Th2 cell differentiation, even under non-polarizing conditions. In mice, HIF2α activation in CD4^+^ T cells promoted intestinal Th2 cell accumulation in the absence of infection, and HIF2α-deficiency impaired CD4^+^ T cell-mediated host immunity to intestinal helminth infection. Our findings identified hypoxia, and the oxygen-regulated transcription factor Hypoxia-Inducible Factor 2α (HIF2α), as key regulators of Th2 cell differentiation and function within the small intestine.

## Introduction

Intestinal helminth infections are among the most prevalent chronic infections worldwide^1^. Helminth infections are often associated with polarized “type 2” immunity, including activation and accumulation of T helper 2 (Th2) cells, type-2 innate lymphoid cells (ILC2), tissue basophils and eosinophils, elevated serum immunoglobulin E (IgE), alternative activation of macrophages and alterations of epithelial differentiation and mucus production that collectively remodel the anatomical site of infection^2^. An analogous cascade of immunological events and tissue remodeling that trigger local tissue pathology occur in allergic diseases, including allergic asthma^3^.

The mechanisms that guide Th2 cell differentiation within barrier tissues remain unclear. However, evidence supporting an instructive role of the tissue microenvironment in establishing protective Th2 cell differentiation and function is provided by changes in chromatin accessibility and/or gene expression following trafficking of Th2 cells from lymph nodes to parenchymal tissues^4,5^. Tissue alarmins, including IL-25, IL-33 and TSLP are key factors produced within barrier tissues that collectively promote type-2 immunity and Th2 cell responses during helminth infection^6,7^. The identity of other cues from the tissue milieu that impact Th2 cell function within barrier tissues remain to be identified.

Hypoxia-inducible factors (HIFs) are key transcription factors that mediate cellular and organismal responses to hypoxia^8^. Consisting of 3 family members, (HIF1α, HIF2α and HIF3α, encoded by *Hif1a, Epas1* and *Hif3a*, respectively), HIFs are post-translationally modified in an oxygen-dependent enzymatic cascade that regulates their stability, nuclear translocation, binding to hypoxia-response elements (HRE) and transcription of hypoxia-inducible genes^8^. Under normoxic conditions HIF proteins are hydroxylated on key proline residues by oxygen-dependent prolylhydroxylase (PHD) enzymes. Recognition of hydroxylated HIF molecules by the ubiquitin ligase Von Hippel-Lindau protein (pVHL) targets HIFs for proteosomal degradation. By contrast, under hypoxic conditions, inactivation of PHD enzymes results in nuclear translocation of HIFs to trigger downstream transcriptional function in concert with HIF1β. Distinct thresholds in oxygen availability trigger activation of discrete HIF members, with HIF1α stabilized at ∼1% O_2_, an oxygen concentration present within ischemic tissues, tumors and the colonic epithelium. By contrast, HIF2α is stabilized below 5% O_2_, an oxygen availability readily found in homeostatic parenchymal tissues, increasingly termed “physoxia”^9^. HIFs are important regulators of activation and effector function of immune cells, including T cells. Most studies have investigated the contributon of HIF1α to effector T cell function^10^. In both CD4^+^ and CD8^+^ T cells, HIF1α can act downstream of both TCR activation and IL-2R-signaling to drive early glycolytic shift following T cell priming^11^. HIF1α is also implicated in the regulation of Th17 and Treg cell differentiation and function^12,13^. By contrast the role of HIF2α in T cell biology is poorly understood. In both macrophages and CD8^+^ T cells activated *in vitro*, HIF2α stabilization can be induced by hypoxia, but notably, also by IL-4 signaling^11,14^. Furthermore, *Epas1* is directly bound by GATA-3 in Th2 cells^15^, suggesting a potential contribution to Th2 cell biology, as was recently described within the lung in a model of experimental asthma^16^.

Here we demonstrate that in the intestine, helminthic infection drives accumulation of small intestinal lamina propria Th2 cells marked by elevated expression of the gene *Epas1*, encoding HIF2α. *In vitro*, activation of HIF2α by culture under a hypoxic atmosphere promotes naïve CD4^+^ T cell differentiation towards a Th2 cell lineage. Strikingly, genetic HIF2α activation also promotes spontaneous Th2 cell differentiation *in vitro* and *in vivo* in the absence of hypoxia or helminthic infection. Finally, deletion of *Epas1*/HIF2α from T cells renders mice susceptible to helminthic reinfection, demonstrating a key role for HIF2α-mediated sensing of tissue physoxia by CD4^+^ T cells and the contribution of this axis to type-2 immunity in the gut.

## Methods and Materials

### Mice

C57BL6/J (WT) were purchased from The Jackson Laboratory. *Epas1*^fl/fl^, HIF2α^LSL^ and *Cd4*^Cre^ mice^17-19^ (The Jackson Laboratory) were intercrossed to generate experimental cohorts and appropriate controls. Mice were infected, and reinfected, with *H. polygyrus* as described^20^. Sex- and age-matched littermate mice (6-12 weeks of age) were randomly assigned to experimental groups. All studies were performed under specific pathogen free conditions, and in accordance with the Institutional Animal Care and Use Committee at Benaroya Research Institute.

### In vitro CD4^+^ T cell cultures

Naïve CD4^+^ T cells were enriched using negative magnetic selection (Miltenyi Biotec) and co-cultured with irradiated splenocytes, agonistic anti-CD3e and anti-CD28 antibodies and either anti-IFN-γ and anti-IL-4 (Th0) or anti-IFN-γ, anti-IL-12 and IL-4 (Th2) in normoxic or hypoxic (4% O_2_) conditions for 96 hours, being split once after 48 hours.

### RNA-seq

Small intestinal lamina propria leukocytes were isolated from naïve and *H. polygyrus* infected (day 30) WT B6 mice and live CD4^+^ T cells were FACS purified prior to capture and processing using V2 10X Genomics 5’ Reagents Kit. *In vitro* activated CD4^+^ T cells were subjected to bulk RNA-seq using Takara SMART-seq v4 Ultra Low Input RNA Kit.

### Cell isolation and flow cytometry

To isolate cells, spleen, lymph node and small intestinal lamina propria cells were isolated and restimulated as previously described^21^. Data were acquired on a Fortessa X-20 or A5 Symphony (BD Biosciences) and analyzed with FlowJo software (Tree Star).

### Statistical analyses

Statistical analyses were performed in Prism (GraphPad Software). *P* values, *≤0.05, **≤0.01, ***≤0.001.

## Results and Discussion

### Epas1 is a transcriptional hallmark of intestinal Th2 cells

We sought to identify novel transcriptional regulators of intestinal Th2 cells elicited by helminth infection. To do so, we used *Heligmosomoides polygyrus*, an intestinal nematode and natural parasite of mice, which establishes a chronic infection of the small intestine and elicits Th2 cell responses in the mesenteric lymph node (MLN) and small intestinal lamina propria (siLP). To identify tissue Th2 cell regulators *in vivo*, we FACS isolated CD4^+^ T cells from the siLP of uninfected WT mice or those infected with *H. polygyrus* (30 d.p.i) and conducted scRNA-seq using the 10X Genomics Chromium system. Post quality control, UMAP identified accumulation of siLP Th2 cells in *H. polygyrus* infected mice **(Fig.1A-C)**, identified by accumulation of cells with elevated *Gata3* expression **(Fig.1D)**. Strikingly, we also identified Th2 cell-specific expression of the Endothelial PAS domain-containing protein 1 (*Epas1*), encoding Hypoxia-inducible factor 2-alpha (HIF2α) **(Fig.1E)**, suggesting this transcription factor may influence siLP Th2 cell differentiation or function during helminth infection. By contrast, family member *Hif1a* was widely expressed by all CD4^+^ T cell subsets present within the siLP regardless of infection status, whereas *Hif3a* was not expressed by any CD4^+^ T cell population analyzed (data not shown). As such, Th2 cells selectively express *Epas1* during intestinal helminthic infection.

**Figure 1.**
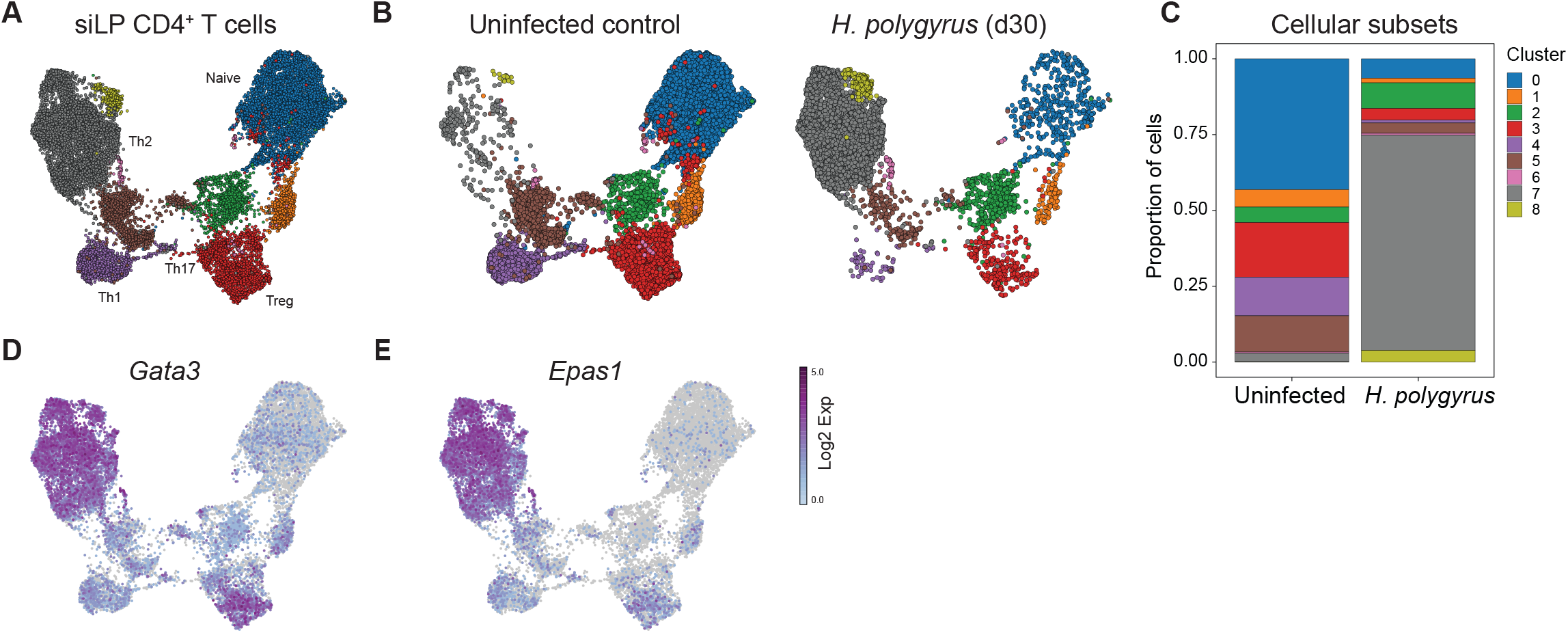
*Epas1* is a transcriptional hallmark of intestinal Th2 cells. CD4^+^ T cells were isolated from naïve WT B6 mice or those infected with *H. polygyrus* (30 d.p.i) and subjected to scRNA-seq. Uniform Manifold Approximation and Projection (UMAP) of (A) cells from both conditions or (B) uninfected mice or *H. polygyrus* infected mice. (C) Relative proportion of cell clusters. (D) *Gata3* and (E) *Epas1* expression among all CD4^+^ T cell clusters from naïve and infected mice combined. Data represent a single experiment with n=2 mice per group.

### Hypoxia promotes Th2 cell differentiation in a Epas1/HIF2α-dependent manner

Given the selective expression of *Epas1* by tissue Th2 cells during helminth infection, and because HIF2α is an oxygen-dependent transcription factor^10^, we hypothesized detection of physiological hypoxia by HIF2α may influence Th2 cell differentiation. To test this, naïve CD4^+^ T cells were cultured under Th0 (anti-IFN-γ, anti-IL-4) and Th2 (anti-IFN-γ, anti-IL-12, IL-4) polarizing conditions in normoxic (5% CO_2_, 20% O_2_) or hypoxic (5% CO_2_, 4% O_2_) atmospheres. As expected, under normoxia, Th2 skewing conditions promoted differentiation of IL-13^+^ Th2 cells **(Fig.2A)**. Strikingly, cultures placed under hypoxia demonstrated a significant increase in the polarization of IL-13^+^Th2 cells **(Fig.2A)**. Thus, physiological hypoxia (4% O_2_) promotes Th2 cell differentiation *in vitro*. Surprisingly, even under non-polarizing Th0 conditions, cells cultured in a hypoxic atmosphere readily differentiated to IL-13^+^ Th2 cells, and this did not readily occur under normoxic conditions **(Fig.2B, C)**. As such, exposure to physiological hypoxia (4% O_2_) promotes spontaneous Th2 cell differentiation *in vitro* and accentuates Th2 cell differentiation under permissive conditions.

**Figure 2.**
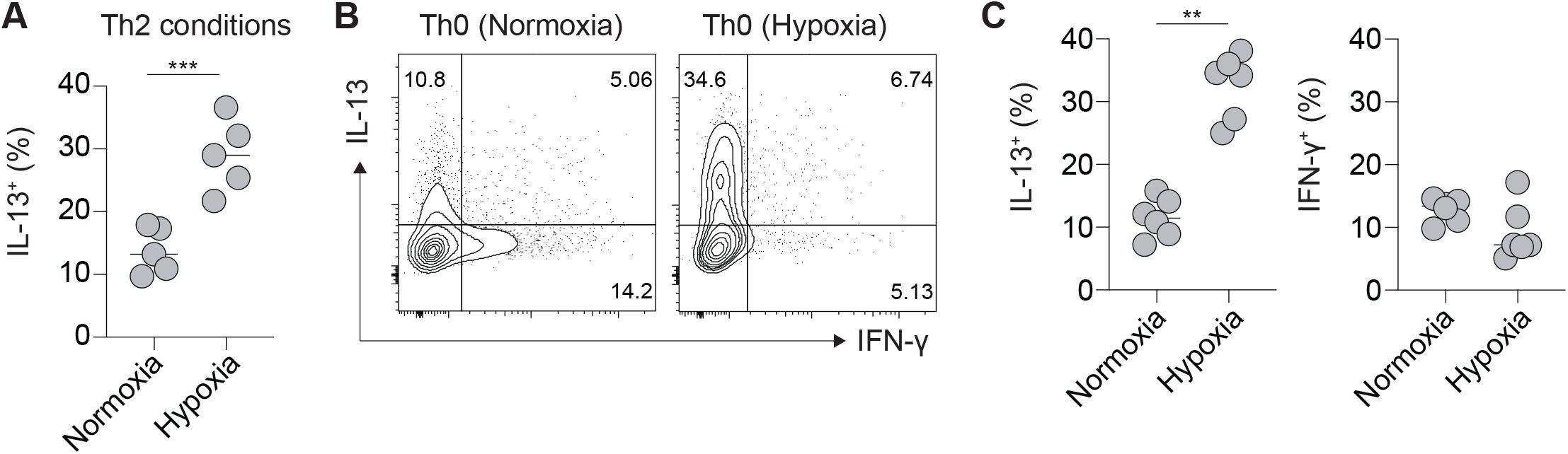
Hypoxia promotes Th2 cell differentiation *in vitro*. (A-C) Naïve CD4^+^ T cells from WT B6 mice were cultured under Th0 or Th2 polarizing conditions under normoxia (20% O_2_) or hypoxia (4% O_2_). (A) Frequencies of IL-13^+^ Th2 cells in cells cultured under Th2 polarizing conditions in normoxia or hypoxia. (B) Representative flow cytometry plots and (C) quantification of IL-13 or IFN-γ production by cells cultured under Th0 conditions under normoxia or hypoxia. Data shown represent one of two independent experiments with up to 5 technical replicates per group. Each dot represents a technical replicate. ** p<0.01, *** p<0.001 (Student’s t-test).

### HIF2α activation is sufficient to promote Th2 cell differentiation

To further understand how selective HIF2α activation influences Th2 cell function, and to distinguish this from the global cellular response to hypoxia, we utilized a constitutively active HIF2α model^18^. Specifically, in HIF2α^LSL^ transgenic mice, within the *Rosa26* locus a floxed stop codon precedes a hemagglutinin (HA)-tagged mutant HIF2α bearing key proline-to-alanine mutations that escape PHD-mediated hydroxylation, generating a non-degradable form of HIF2α that is transcriptionally active under normoxia^18^. We intercrossed HIF2α^LSL^ mice with *Cd4*^Cre^ mice to drive constitutive HIF2α activation in CD4^+^ and CD8^+^ T cells (HIF2α^CD4-ON^ mice). Notably, when cultured *in vitro*, naïve CD4^+^ T cells with constitutive HIF2α activation demonstrated spontaneous Th2 cell differentiation under normoxia **(Fig.3A,B)**, indicating that HIF2α activation alone mimics hypoxia and is sufficient to promote IL-13^+^ Th2 cell differentiation. To determine whether transcriptional regulation of Th2 cell differentiation by hypoxia and HIF2α extend beyond production of IL-13, we conducted bulk RNA-seq on WT naïve CD4^+^ T cells cultured under Th0 conditions in normoxia and hypoxia, as well as on naïve CD4^+^ T cells from HIF2α^LSL^ and HIF2α^CD4-ON^ mice cultured under Th0 conditions in normoxia. Differential gene expression analysis identified a high concordance in transcriptional signature of Th2 cells elicited by culture in hypoxic conditions, or through genetic activation of HIF2α, including *Il5, Il10* and *Zeb2* **(Fig. 3C)**.

**Figure 3.**
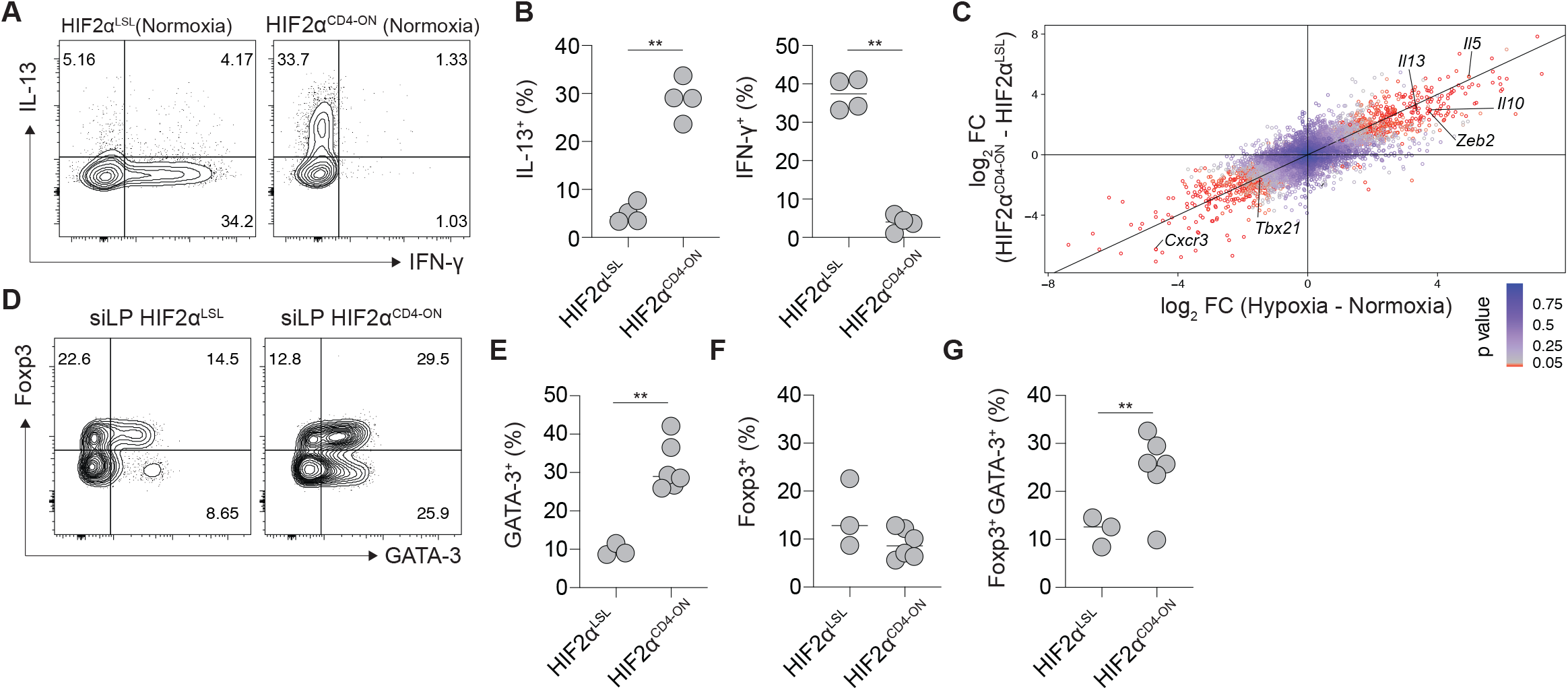
Genetic HIF2α activation promotes Th2 cell differentiation *in vitro* and *in vivo*. (A-B) Naïve CD4^+^ T cells from HIF2α^LSL^ and HIF2α^CD4-ON^ mice were cultured without skewing under normoxia. Representative flow cytometry plots (A) and quantification (B) of IL-13 and IFN-γ production. (C) Differential gene expression analyses of bulk RNA-seq of naïve CD4^+^ T cells from WT mice cultured under Th0 conditions in normoxia and hypoxia, and naïve CD4^+^ T cells from HIF2α^LSL^ and HIF2α^CD4-ON^ mice cultured under Th0 conditions in normoxia. (D-G) CD4^+^ T cell subsets present in the small intestinal lamina propria of naïve HIF2α^LSL^ and HIF2α^CD4-ON^ mice were assessed by flow cytometry. (D) Representative flow cytometry plots and (E-G) quantification of cells expressing GATA-3, and/or Foxp3. Data shown represent one of two independent experiments with 3-5 technical or biological replicates, except for C, which was conducted once with 3 technical replicates. ** p<0.01 (Student’s t-test).

To determine how HIF2α activation influences CD4^+^ T cell differentiation *in vivo*, we aged experimental cohorts of HIF2α^CD4-ON^ and HIF2α^LSL^ control littermates. While HIF2α^LSL^ mice developed normally, older HIF2α^CD4-ON^ mice developed lymphadenopathy and splenomegaly (data not shown), indicating that perturbation of normal oxygen sensing through genetic activation of HIF2α is sufficient to perturb immune homeostasis *in vivo*. Notably, secondary lymphoid organs of HIF2α^CD4-ON^ mice demonstrated accumulation of activated CD4^+^ and CD8^+^ T cells (data not shown), but most strikingly, HIF2α^CD4-ON^ mice demonstrated a prominent accumulation of both GATA-3^+^ Th2 cells **(Fig.3D,E)**, as well as GATA-3^+^Foxp3^+^ Treg cells **(Fig.3D,F,G)** within barrier tissues, including the gastrointestinal tract, that was not observed in HIF2α^LSL^ controls **(Fig.3D-G)**. Whether accumulation of GATA-3^+^Foxp3^+^ Treg cells occurs in response to Th2 cell accumulation or is due to a cell-intrinsic role of HIF2α transgene activation in Foxp3^+^ Treg requires further investigation. However, even in the absence of helminth infection, constitutive activation of HIF2α in CD4^+^ T cells is sufficient to promote accumulation of Th2 cells within tissue microenvironments. Thus HIF2α-dependent oxygen sensing represents a key checkpoint in Th2 cell differentiation.

### Th2 cell-mediated protection from helminth infection requires HIF2a

Whereas genetic activation of HIF2α is sufficient to promote Th2 cell differentiation within the siLP, we sought to determine the necessity of this factor in Th2 cell mediated host protection. While *H. polygyrus* establishes a chronic primary infection, following drug-mediated worm clearance, CD4^+^ T cells provide sterilizing protection against reinfection^20^. We used this setting to investigate the necessity for HIF2α in protective CD4^+^ T cell function *in vivo*. Specifically, we generated mice with T cell-specific HIF2α deficiency, by intercrossing *Epas1*^fl/fl^ and *Cd4*^Cre^ mice. *Epas1*^fl/fl^ and *Epas1*^ΔCD4^ littermates were infected with *H. polygyrus*, and 14 days post infection were dewormed using pyrantel pamoate, before being reinfected with *H. polygyrus* 28 days later **(Fig. 4A)**. In control *Epas1*^fl/fl^ mice, CD4^+^ T cell-mediated protection was intact, whereas *Epas1*^ΔCD4^ mice displayed increased susceptibility to reinfection, with increased worm burden and fecundity upon reinfection **(Fig. 4B,C)**. As such, T cell-intrinsic expression of *Epas1* is required for protective immunity to helminth infection within the siLP.

**Figure 4.**
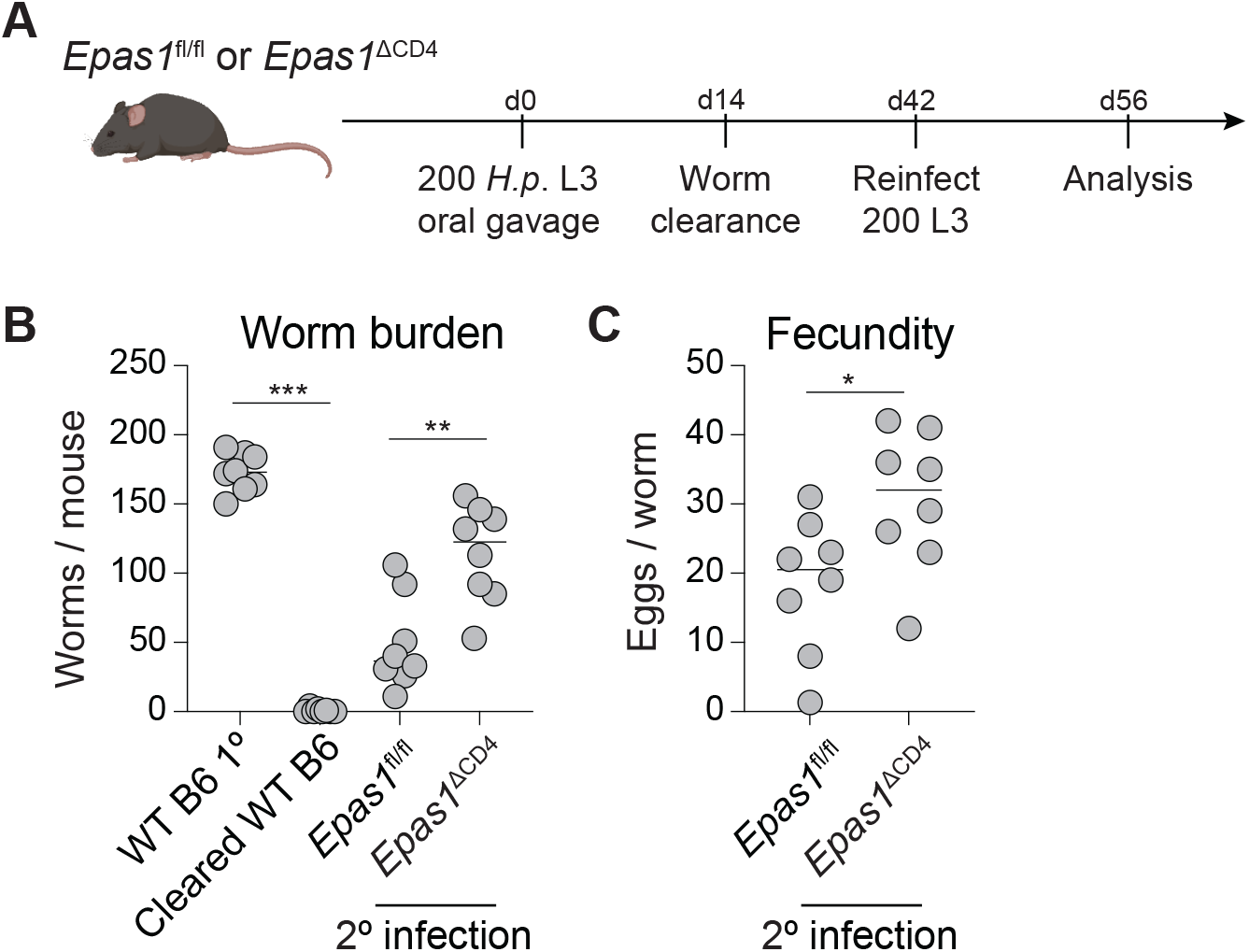
T cell-intrinsic expression of HIF2α is required for host immunity to intestinal helminth infection. (A) Experimental schematic for determining CD4^+^ T cell mediated host immunity in *Epas1*^fl/fl^ and *Epas1*^ΔCD4^ mice. (B) Worm burden and (C) worm fecundity from *Epas1*^fl/fl^ and *Epas1*^ΔCD4^ mice reinfected with *H. polygyrus*. Data shown is pooled from 2 independent experiments with n=3-5 mice per group. * p<0.05, ** p<0.01, *** p<0.001. (one-way ANOVA with Holm-Šidák multiple-comparison test (B), or Student’s t-test (C)).

In summary, our work provides support that HIF2α, through sensing of reduced oxygen availability, acts in a cell-intrinsic manner to promote Th2 cell differentiation and acts to promote CD4^+^ T cell mediated immunity to gastrointestinal helminth infection. Future studies will be required to determine how HIF2α impacts Th2 cell differentiation, retention and function in the helminth infected intestine.

## Declaration of Interests

The authors declare no competing financial interests.

## Lead contact and materials availability

Sequencing data is deposited under GEO: (Upload in progress).

All correspondence and material requests should be addressed to O.J.H.

## Acknowledgements

We thank Jakob von Moltke (University of Washington) for provision of *H. polygyrus* L3 larvae. We thank staff of Benaroya Research Institute Cell and Tissue Analysis and Genomics Cores, particularly Adam Wojno, Vivian Gersuk, Kimberley O’Brien and Quynh-Anh Nguyen, as well as staff of the Benaroya Research Institute vivarium. We thank the M.J. Murdock Charitable Trust for support of flow cytometry, histology, imaging and genomics resources at Benaroya Research Institute.

## Funding

This work was supported by Benaroya Research Institute, the Gut Immunity Program at Benaroya Research Institute. S.V is a Washington Research Foundation Post-doctoral Fellow, T.M.O. is the recipient of a T32 CMB training grant (1T32GM136534), and a National Science Foundation GRFP fellowship (DGE-2140004). O.J.H was supported by NIH (R01AI158624).

## Notes

### Competing Interest Statement

The authors have declared no competing interest.

